# Surface sensing and metabolic signals drive feedback regulation of long polar fimbriae in enterohemorrhagic *Escherichia coli* O157

**DOI:** 10.64898/2026.10.14.759193

**Authors:** S Dutta, W Galdamez, MA Ramirez, AM Krachler

## Abstract

Bacterial colonization requires coordinated regulation of adhesion structures in response to environmental conditions. While metabolic regulation of adhesin expression is well established, how surface sensing contributes to this regulation remains less understood. Here, we investigated the regulation and function of long polar fimbriae (Lpf) in enterohemorrhagic *Escherichia coli* O157. We show that the *lpf1* and *lpf2* promoters integrate metabolic and surface-associated signals through distinct regulatory regions. Bile induces both promoters independently of distal regulatory sequences, whereas these regions repress basal transcription and are required for induction following surface association. Surface-dependent activation of both *lpf* promoters is reduced in mutants lacking the major fimbrial subunits LpfA1 and LpfA2, but is unaffected by deletion of the distal tip adhesins. Lpf structures also promote surface-dependent activation of *ler*, a positive regulator of *lpf* expression, supporting a regulatory feedback circuit linking surface sensing to *lpf* transcription. Loss of the major fimbrial subunits also impairs colonization in zebrafish, whereas deletion of the tip adhesins does not under the conditions tested. Together, these findings show that Lpf integrate environmental signals through surface-dependent regulatory feedback and identify an additional role for the fimbrial structure during colonization that does not require the tip adhesins.

## INTRODUCTION

Enterohemorrhagic *Escherichia coli* (EHEC) is an enteric pathogen that colonizes the intestinal tract and can cause diarrheal disease, hemorrhagic colitis, and hemolytic uremic syndrome. During infection, bacteria encounter heterogeneous environments characterized by fluid flow, surface contact, nutrient gradients, and host-derived stresses. These conditions vary across spatial and temporal scales, requiring coordinated regulation of virulence-associated genes during colonization (Dutta *et al*., 2026). The locus of enterocyte effacement (LEE) encodes a type III secretion system and associated regulators required for EHEC virulence. The LEE-encoded regulator Ler functions as a central activator of this system and controls expression of multiple virulence genes. Ler activity is modulated by many transcriptional regulators, including the transcriptional activator GrlA (Barba *et al*., 2005), and the histone-like protein H-NS, which acts as a repressor (Laaberki *et al*., 2006). Previous work has shown that Ler induction and LEE expression are responsive to surface-associated conditions and mechanical stimulation, linking host attachment to activation of virulence gene expression via GrlA (Sirisaengtaksin *et al*., 2020, Alsharif *et al*., 2015). Thus, surface sensing provides an environmental input into the regulatory networks controlling EHEC virulence.

Long polar fimbriae (Lpf) are chaperone–usher fimbriae encoded by enterohemorrhagic *E. coli* (EHEC), other pathogenic *E. coli* strains, and diverse enteric pathogens, including *Salmonella* species (Baumler & Heffron, 1995, Torres *et al*., 2002, Newton *et al*., 2004). The *lpf* operons encode the major fimbrial subunit (LpfA), periplasmic chaperone, outer membrane usher, and distal subunits required for fimbrial assembly (Torres *et al*., 2002). Lpf contribute to colonization *in vivo* (Jordan *et al*., 2004), and the distal tip adhesins, LpfE1 and LpfD′2 for Lpf1 and Lpf2, respectively, have been implicated in receptor-mediated adhesion (Coppens *et al*., 2015). Chaperone–usher fimbrial shafts can also have important biomechanical properties, including the ability to extend under force and dissipate mechanical stress (Spaulding *et al*., 2018). Lpf have been associated with adherence to intestinal tissues, including Peyer’s patches and M cells, and have been shown to bind extracellular matrix components (Baumler *et al*., 1996, Farfan *et al*., 2011, Torres *et al*., 2008). The presence of *lpf* genes correlates with pathogenic strains, and Lpf contribute to intestinal persistence in animal models (Cordonnier *et al*., 2017, Torres *et al*., 2004, Torres *et al*., 2009). In addition, Lpf have been implicated in modulation of host immune responses (Vergara *et al*., 2014). Despite these observations, the mechanisms by which Lpf contribute to colonization remain incompletely defined. Expression of *lpf* operons is controlled by multiple regulatory inputs. H-NS represses transcription under non-permissive conditions, while Ler can relieve this repression through direct interaction with *lpf* promoters (Torres *et al*., 2007a, Rojas-Lopez *et al*., 2011). Environmental factors including growth phase, osmolarity, pH, iron availability, and host-associated metabolites such as bile have been shown to influence *lpf* expression (Arenas-Hernandez *et al*., 2014, Chassaing *et al*., 2013, Hu *et al*., 2015). These observations indicate that Lpf are subject to complex regulation, but how different environmental signals are integrated at the *lpf* promoters remains unclear.

EHEC encodes multiple fimbrial systems, and regulatory cross-talk between adhesion loci has been described in enteric bacteria (Gonyar *et al*., 2024, Lloyd *et al*., 2012). Expression of individual fimbriae can influence the regulation of others, and coordinated control of adhesion systems contributes to adaptation during colonization. Recent studies have also shown that fimbrial loci can affect global gene expression, indicating roles beyond direct adhesion (Gonyar *et al*., 2024). Whether Lpf themselves contribute to regulatory responses following surface association, and whether such regulation is linked to their expression, is not known. Chaperone–usher fimbriae are typically described as adhesive structures in which the distal subunit mediates receptor binding and the fimbrial shaft provides structural support. However, the contribution of these different structural components to functions beyond adhesion is less well understood. In particular, whether the assembled fimbrial structure can influence transcriptional regulation following surface association has not been established.

Here, we examined how surface-associated and metabolic signals regulate *lpf* expression and Lpf-dependent colonization. We show that *lpf1* and *lpf2* integrate surface association and bile through distinct promoter regions, and that surface-dependent induction requires the Lpf fimbrial structure but not the distal tip adhesins. Lpf also promote surface-dependent activation of Ler, supporting a feedback circuit that enhances *lpf* expression. Distal H-NS–associated promoter regions constrain basal expression and enable surface responsiveness, whereas bile responsiveness is retained proximally. Together, these findings reveal a regulatory role for Lpf in surface sensing and colonization that extends beyond tip-mediated adhesion.

## RESULTS

### Distal regulatory regions constrain basal *lpf* promoter activity but are dispensable for bile induction

To define the regulatory architecture of *lpf* expression, we analyzed *lpf1* and *lpf2* promoter activity using *lacZ* and *gfp* transcriptional reporters. For each operon, we generated a “long” reporter encompassing the full intergenic regulatory region (350 bp for *lpf1* and 310 bp for *lpf2*, termed P*lpf1*_L_ and P*lpf2*_L_, respectively) and a truncated “short” reporter containing the proximal 200 bp upstream of the translational start site (P*lpf1*_S_ and P*lpf2*_S_, respectively). The distal regions removed from the short constructs contain previously identified regulatory sites, including those associated with the global repressor H-NS and the anti-silencer Ler, as well as Fur for *lpf2* (Torres *et al*., 2007b, Torres *et al*., 2007a).

In planktonic cultures, all four reporters showed growth phase-independent activity in DMEM (Fig. S1), and all subsequent experiments were carried out on mid-log phase cultures. Basal promoter activity was approximately 3-fold higher for the P*lpf2*_L_ promoter compared to P*lpf1*_L_ (Fig. 1A). Upon addition of 0.15% bile, both *lpf1* and *lpf2* promoters were moderately but significantly induced (approximately 1.5-fold for P*lpf1*_L_ and 1.3-fold for P*lpf2*_L_; Fig. 1A). Truncation of the distal regulatory regions increased basal activity of both promoters (11-fold for P*lpf1*_*S*_ and 7-fold for P*lpf2*_*S*_), and they both remained inducible by bile (Fig. 1B). We obtained similar results with *gfp*-based fluorescence reporters (Fig. 1C, D). These findings are consistent with distal repression of both *lpf* promoters, and indicate that bile responsiveness is retained within the proximal 200-bp regions rather than requiring relief of distal repression. Together, these results indicate that distal regulatory elements maintain a low basal state, whereas the proximal promoters retain responsiveness to host-derived metabolic cues such as bile. The persistence of bile induction after removal of the distal regions further suggests that bile and other environmental signals acting through the distal regulatory regions are integrated through distinct features of the *lpf* promoters.

**Fig. 1.**
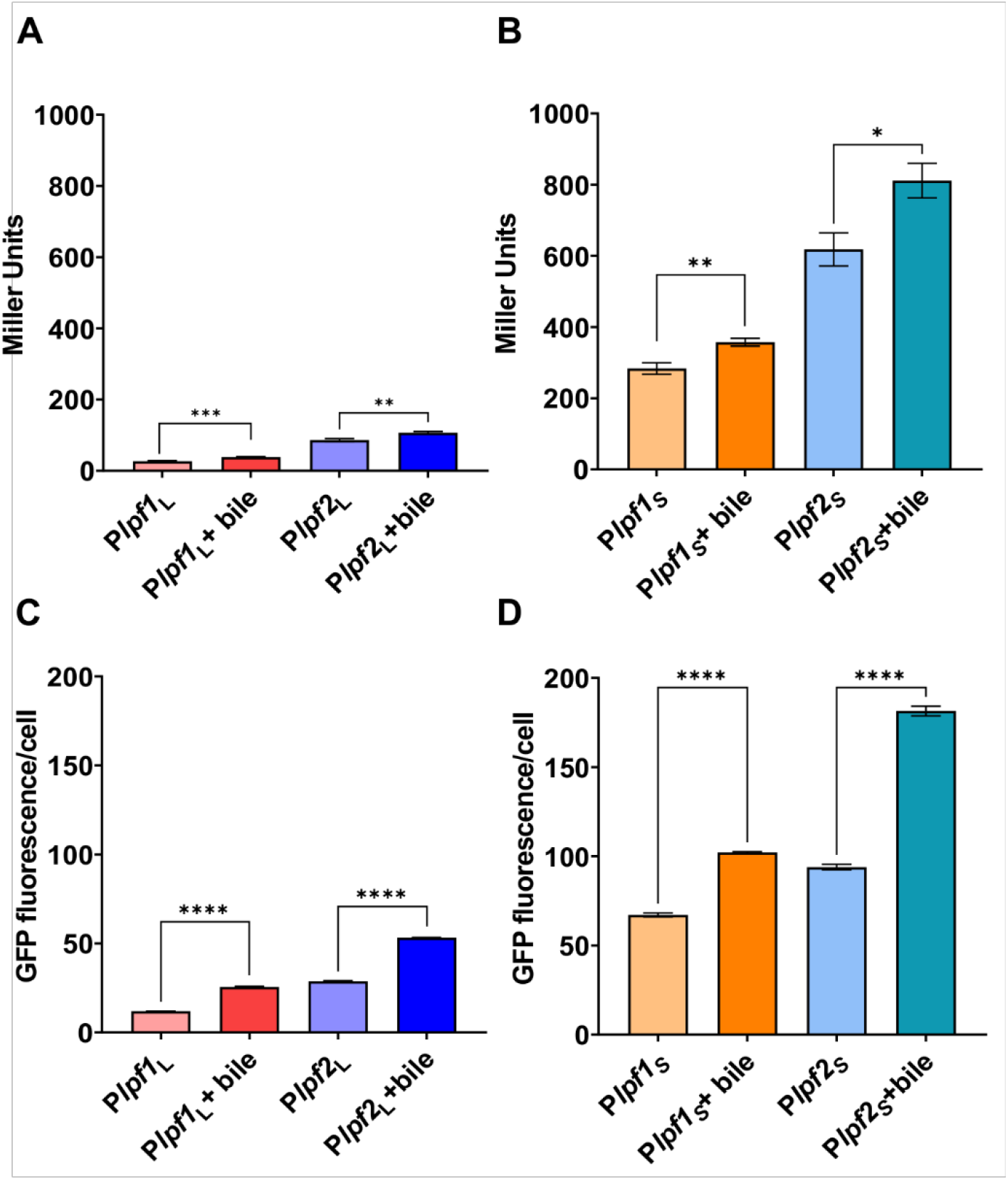
Distal regulatory regions constrain basal *lpf* promoter activity but are dispensable for bile induction. **(A, B)** β-galactosidase activity of transcriptional *lacZ* fusions to the long promoters, P*lpf1*_L_ and P*lpf2*_L_ **(A)**, and short promoters, P*lpf1*_S_ and P*lpf2*_S_ **(B)**, in wild-type EHEC 86-24 grown planktonically in DMEM with or without 0.15% bile. Data represent mean ± SEM from three independent biological replicates (n = 3). Statistical significance was determined by one-way ANOVA with Sidák’s multiple-comparison test (*p<0.05; **p<0.01; ***p<0.001). **(C, D)** GFP fluorescence from transcriptional *gfp* fusions to P*lpf1*_L_ and P*lpf2*_L_ **(C)** and P*lpf1*_S_ and P*lpf2*_S_ **(D)** under the same growth conditions. Data represent mean ± SEM of normalized GFP fluorescence per cell from at least 150 cells per condition across three independent biological replicates (n = 3). Statistical significance was determined by one-way ANOVA with Sidák’s multiple-comparison test (****p<0.0001).

### *lpf* promoter architecture integrates surface-associated and metabolic cues

We have previously shown that surface sensing regulates EHEC virulence gene expression (Alsharif *et al*., 2015, Sirisaengtaksin *et al*., 2020). We therefore tested whether the *lpf* promoters also respond to surface association, using P*lpf1*_L_:*gfp* and P*lpf2*_L_:*gfp* transcriptional reporters in wild-type EHEC. The GFP and β-galactosidase reporters showed comparable patterns of basal activity and bile-dependent induction, with approximately 2-fold induction of the GFP reporters following addition of bile (Fig. 1C, D). qRT-PCR measurements of *lpf1* and *lpf2* transcript levels validated bile induction across genes in both operons (Fig. S2). Compared with planktonic cells, surface-associated cells showed approximately 3-fold and 2-fold increases in mean GFP fluorescence from P*lpf1*_L_ and P*lpf2*_L_, respectively (Fig. 2A-C).

**Fig. 2.**
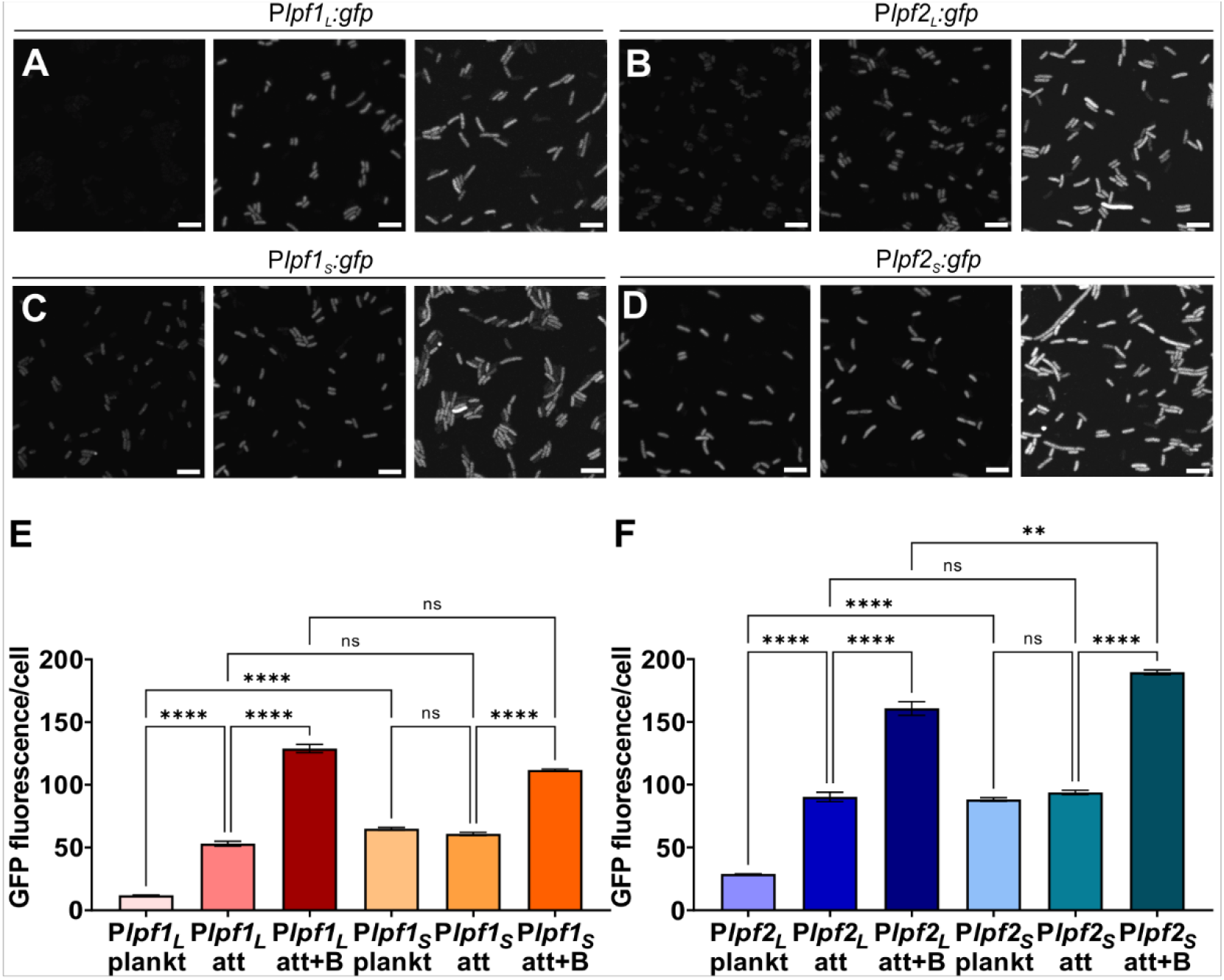
*lpf* promoter architecture integrates surface-associated and metabolic cues. **(A–D)** Representative fluorescence images of P*lpf1*_L_:*gfp* **(A)**, P*lpf2*_*L*_:*gfp* **(B)**, P*lpf1*_S_:*gfp* **(C)**, and P*lpf2*_S_:*gfp* **(D)** transcriptional reporters in wild-type EHEC 86-24 grown planktonically, surface-associated, or surface-associated in the presence of 0.15% bile. Scale bars, 5 µm. **(E–F)** Background-corrected GFP fluorescence intensity per cell, normalized to cell area, for P*lpf1*_L_:*gfp* and P*lpf1*_S_:*gfp* **(E)** and P*lpf2*_L_:*gfp* and P*lpf2*_S_:*gfp* **(F)** under the indicated conditions. Data represent mean ± SEM from three independent biological replicates, with ≥150 cells analyzed per condition per experiment (n = 3). Statistical significance was determined using the Kruskal–Wallis test followed by Dunn’s multiple-comparisons test (ns, p≥0.05; **p<0.01; ****p<0.0001).

We next asked whether this surface-associated response depended on promoter architecture by comparing the long and short promoter constructs for both *lpf1* and *lpf2*. In planktonic cells, the short promoters exhibited substantially higher basal activity than their long counterparts (Fig. 1C), consistent with repression mediated by the distal regulatory regions. In contrast to P*lpf1*_L_ and P*lpf2*_L_, which were induced following surface association (Fig. 2A-C), P*lpf1*_S_ and P*lpf2*_S_ remained at their elevated activity and showed little or no further induction (Fig. 2D-F). Thus, removal of the distal regulatory regions increases basal transcription but eliminates surface-dependent induction.

We then examined how surface-associated and host-derived metabolic signals are integrated at these promoters. Addition of 0.15% bile increased promoter activity under planktonic conditions for both long and short constructs, confirming that bile responsiveness is retained within the proximal promoter regions (Fig. 1C). Bile further enhanced *lpf* expression in surface-associated cells by approximately 2-fold for all four reporters compared with untreated surface-associated cells (Fig. 2), similar to the magnitude of bile-dependent induction observed in planktonic cultures (Fig. 1C). Thus, bile responsiveness is maintained following surface association and does not require the distal regulatory regions.

Together, these results indicate that the distal regulatory regions of the *lpf* promoters both constrain basal transcription and are required for surface-dependent induction. P*lpf1*_L_ and P*lpf2*_L_ therefore transition from low activity in planktonic cells to increased activity following surface association, whereas P*lpf1*_S_ and P*lpf2*_S_ exhibit elevated basal activity and little further response to surface contact. In contrast, bile responsiveness is retained after removal of the distal regions, suggesting that metabolic and surface-associated inputs act through separable features of *lpf* promoter architecture.

### Surface-dependent induction of *lpf* promoters is modulated by Lpf structures but is independent of tip adhesins

To determine whether Lpf structures contribute to surface-dependent regulation, we compared *lpf* promoter activity in wild-type EHEC and mutants lacking major fimbrial or distal tip subunits. Since *lpf* genes are organized into an operon and transcribed as a single polycistronic mRNA (Torres *et al*., 2002), strains lacking either major fimbrial subunits for Lpf1 (Δ*lpfA1*) or Lpf2 (Δ*lpfA2*), as well as the double mutant (Δ*lpfA1*Δ*lpfA2*) lack both Lpf1 and Lpf2. Under planktonic conditions, P*lpf1*_L_ and P*lpf2*_L_ promoter activity was low, and deletion of *lpfA1, lpfA2*, both major fimbrial genes together, or either tip adhesin gene (*lpfE1* or *lpfD’2*) did not significantly alter basal activity of either promoter (Fig. 3A, C).

**Fig. 3.**
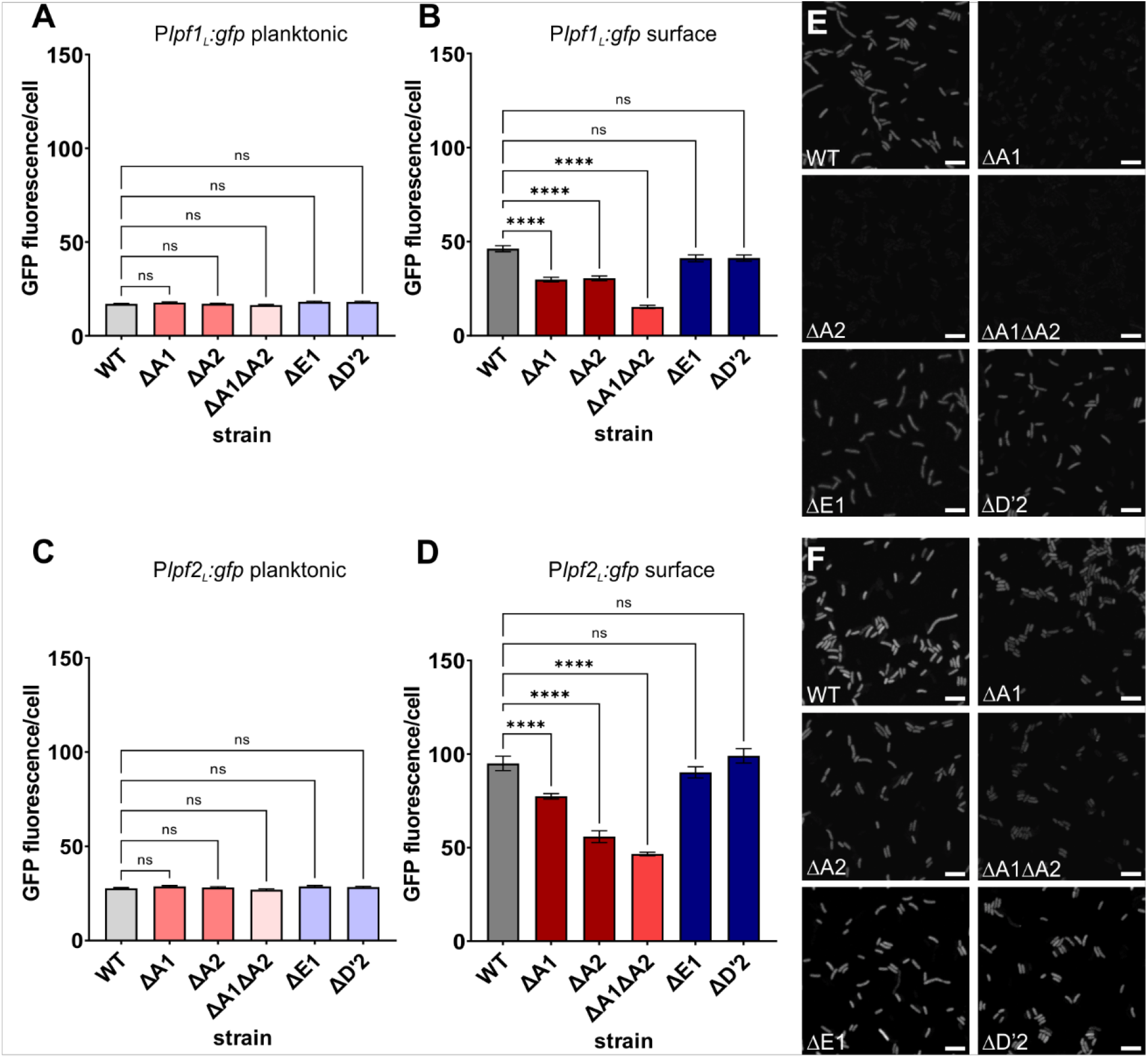
Surface-dependent induction of *lpf* promoters is modulated by Lpf structures but is independent of tip adhesins. **(A–D)** Background-corrected GFP fluorescence intensity per cell, normalized to cell area, for *lpf*:*gfp* transcriptional reporters in wild-type EHEC and *lpf* mutants Δ*lpfA1*, Δ*lpfA2*, Δ*lpfA1*Δ*lpfA2*, Δ*lpfE1*, and Δ*lpfD′2*. P*lpf1*_L_:*gfp* in planktonic **(A)** and surface-associated **(B)** cells and P*lpf2*_L_:*gfp* in planktonic **(C)** and surface-associated **(D)** cells. Data represent mean ± SEM from three independent biological replicates, with ≥150 cells analyzed per condition per experiment (n = 3). Statistical significance was determined using one-way ANOVA with Dunnett’s multiple-comparisons test (ns, p≥0.05; **p<0.01; **p<0.0001). **(E–F)** Representative fluorescence images of surface-associated wild-type and *lpf* mutant cells carrying *Plpf1*_L_:*gfp* **(E)** or P*lpf2*_L_:*gfp* **(F)**. Scale bars, 5 µm.

In wild-type cells, surface association increased both P*lpf1*_L_ and P*lpf2*_L_ activity by approximately 3-fold (Fig. 3B, D). Surface-dependent induction of both promoters was reduced in the Δ*lpfA1* and Δ*lpfA2* single mutants, while the Δ*lpfA1*Δ*lpfA2* double mutant showed the strongest phenotype, with a marked loss of surface-dependent induction, particularly for P*lpf1*_L_ (Fig. 3A-D). Importantly, the reduced response observed in the fimbrial mutants was not attributable to impaired surface association, as cells were immobilized on agarose pads under conditions in which surface association does not depend on Lpf. In contrast, deletion of either distal tip adhesin gene, *lpfE1* or *lpfD’2*, did not substantially affect surface-dependent induction of P*lpf1*_L_ or P*lpf2*_L_, with both tip mutants showing responses similar to wild type (Fig. 3E, F).

Together, these results indicate that Lpf structures are not necessary for surface-dependent induction, they contribute to surface-dependent activation of their own promoters, consistent with positive feedback on *lpf* expression. This feedback requires the major fimbrial subunits but not the distal tip adhesins, distinguishing the contribution of the fimbrial structure from tip-mediated adhesion.

### Ler contributes to feedback between Lpf surface sensing and *lpf* expression

Because Ler is a positive regulator of *lpf* expression (Torres *et al*., 2007a) and we previously found that *ler* itself is induced following surface association (Alsharif *et al*., 2015), we asked whether Lpf structures influence surface-dependent *ler* activation. We measured *ler* promoter activity using a P*ler:gfp* transcriptional reporter in wild-type EHEC and the *lpf* mutants. Under planktonic conditions, deletion of the major fimbrial genes *lpfA1* or *lpfA2*, alone or in combination, or either distal tip adhesin gene, *lpfE1* or *lpfD’2*, did not significantly alter *ler* promoter activity compared with wild type (Fig. 4A).

**Fig. 4.**
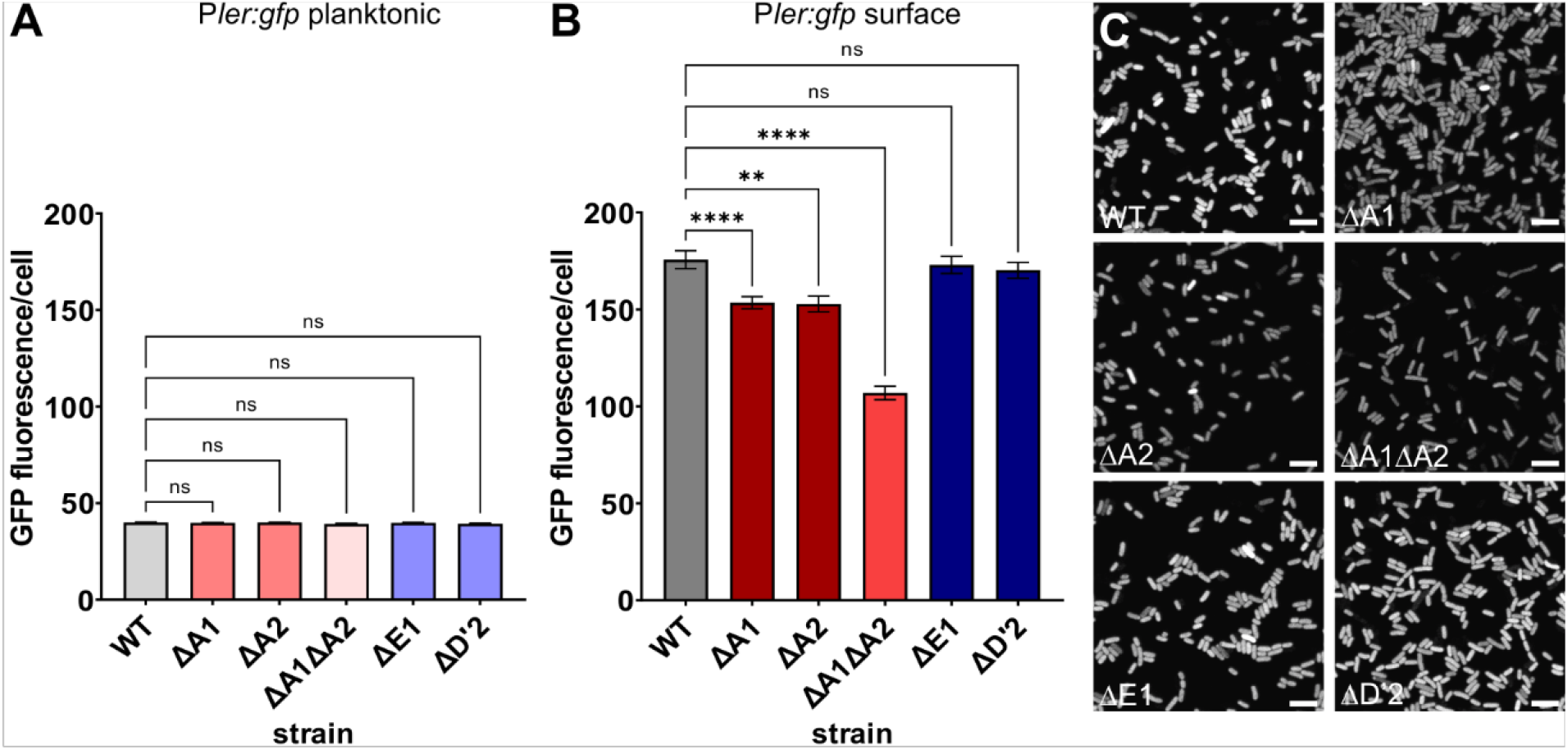
Ler contributes to feedback between Lpf surface sensing and *lpf* expression. **(A–B)** Background-corrected GFP fluorescence intensity per cell, normalized to cell area, from a P*ler:gfp* transcriptional reporter in wild-type EHEC 86-24 and Δ*lpfA1*, Δ*lpfA2*, Δ*lpfA1*Δ*lpfA2*, Δ*lpfE1*, and Δ*lpfD′2* mutants grown planktonically **(A**) or surface-associated **(B)**. Data represent mean ± SEM from three independent biological replicates, with ≥150 cells analyzed per condition per experiment (n = 3). Statistical significance was determined using one-way ANOVA with Dunnett’s multiple-comparisons test, comparing each mutant with wild type (ns, p≥0.05; **p<0.01; ****p<0.0001). **(C)** Representative fluorescence images of surface-associated wild-type and *lpf* mutant cells carrying the P*ler:gfp* reporter. Scale bars, 5 µm.

Following surface association, *ler* promoter activity increased approximately 4-fold in wild-type cells compared to cells grown planktonically in DMEM (Fig. 4A, B). This surface-dependent induction was reduced in Δ*lpfA1* and Δ*lpfA2* cells and was further reduced in the Δ*lpfA1*Δ*lpfA2* double mutant, closely paralleling the effects observed for the *lpf1* and *lpf2* promoters. In contrast, deletion of *lpfE1* or *lpfD’2* did not substantially impair surface-dependent *ler* activation, with both tip mutants resembling wild type (Fig. 4B). Representative images illustrate the differences in *ler* promoter activity between surface-associated wild-type and *lpf* mutant cells (Fig. 4C).

Thus, the same Lpf structural features that promote surface-dependent *lpf* expression also contribute to surface-dependent activation of *ler*. Given the established role of Ler as a positive regulator of *lpf* expression, these findings support a regulatory feedback circuit in which Lpf structures promote surface-dependent *ler* activation, which in turn enhances *lpf* expression. Of note, *ler* induction in response to surface-association was still significant compared to planktonic cells in Δ*lpfA1*Δ*lpfA2*, indicating that even though Lpf contribute to surface-mediated induction, they are not completely essential, and additional elements likely contribute to mechanosensing.

### Lpf-dependent colonization requires the fimbrial shaft but not the tip adhesins

To assess the functional relevance of Lpf *in vivo*, we quantified intestinal colonization in a zebrafish model of EHEC colonization (Stones *et al*.). Even though all strains grew similarly in culture (Fig. S4), deletion of the major fimbrial subunits reduced intestinal bacterial burden, with the Δ*lpfA1*Δ*lpfA2* double mutant showing the strongest colonization defect (Fig. 5A). In contrast, deletion of either distal tip adhesin, *lpfE1* or *lpfD’2*, did not significantly alter bacterial burden compared with wild type (Fig. 5A). Thus, under these conditions, Lpf-dependent colonization requires the major fimbrial subunits but not the distal tip adhesins. Fluorescence imaging of infected animals was consistent with the CFU measurements, with reduced intestinal bacterial signal in the major fimbrial mutants but not the tip adhesin mutants (Fig. 5B). Quantification of intestinal fluorescence showed the same overall pattern (Fig. 5C).

**Fig. 5.**
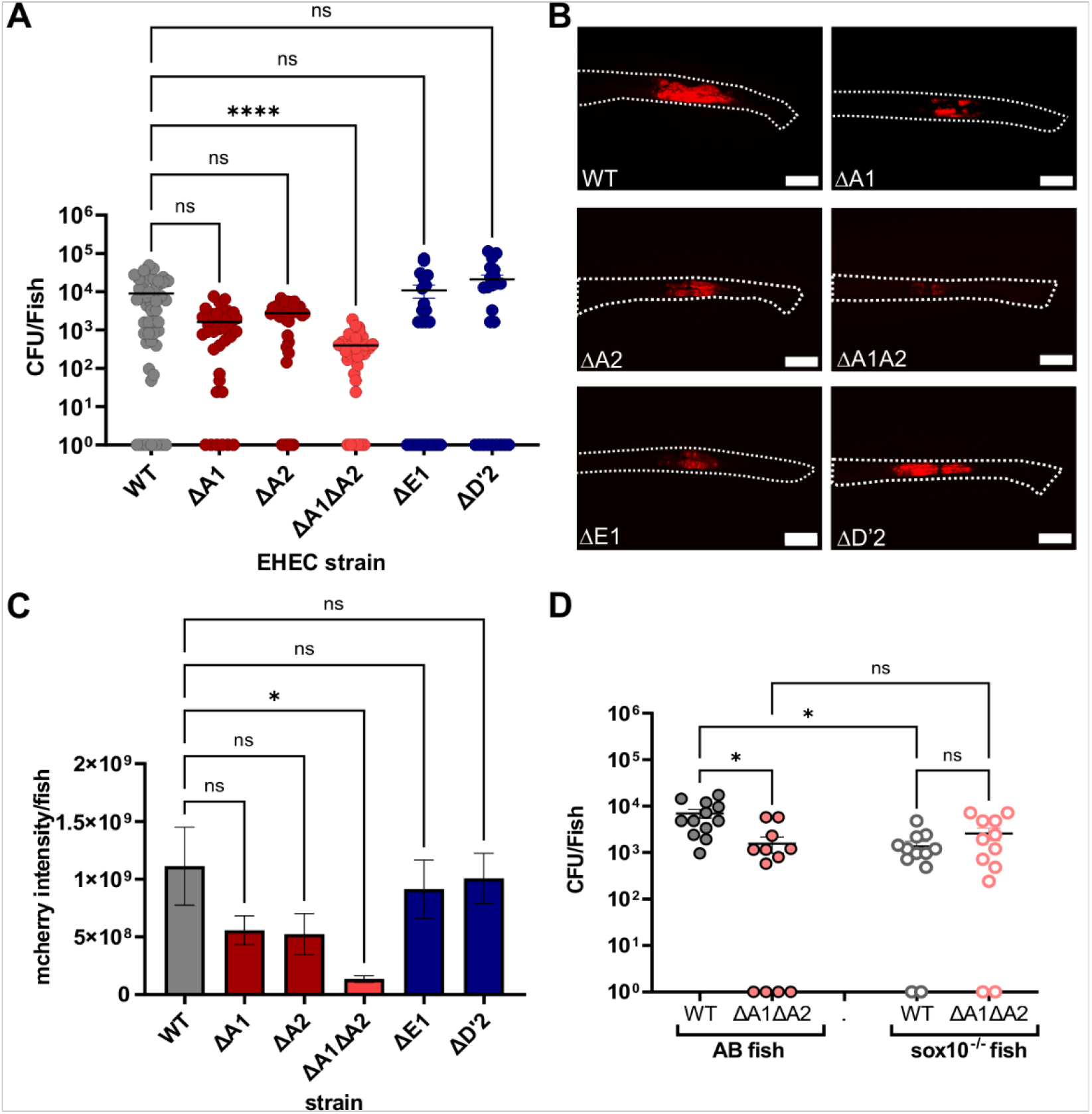
Lpf-dependent colonization requires the major fimbrial subunits but not the tip adhesins. **(A)** Intestinal colonization of zebrafish larvae infected with wild-type EHEC, Δ*lpfA1*, Δ*lpfA2*, Δ*lpfA1*Δ*lpfA2*, Δ*lpfE1*, or Δ*lpfD′2*. Bacterial burden is shown as CFU/fish. Dots represent individual larvae; data are mean ± SEM from ≥3 independent experiments (≥24 larvae/strain). Statistical significance was determined by the Kruskal–Wallis test with Dunn’s multiple-comparisons test (ns, p≥0.05; ****p<0.0001). Samples with undetectable bacterial burden were assigned a value of 1 for visualization and analysis. **(B)** Representative images of larvae infected with mCherry-expressing wild-type or *lpf* mutants; intestines outlined (white dashed lines). Scale bars, 100 µm. **(C)** Intestinal fluorescence intensity of infected larvae. Data are mean ± SEM (n=3). Statistical significance was determined by one-way ANOVA with Dunnett’s multiple-comparisons test (ns, p≥0.05; *p<0.05). **(D)** Intestinal colonization of wild-type AB and *sox10*^*−/−*^ larvae infected with wild-type EHEC or Δ*lpfA1*Δ*lpfA2*. Bacterial burden is shown as CFU/fish; dots represent individual larvae. Data are mean ± SEM (n≥3 independent experiments and ≥12 larvae/condition). Statistical significance was determined by the Kruskal–Wallis test with Dunn’s multiple-comparisons test (ns, p≥0.05; *p<0.05).

To confirm that the phenotypes associated with loss of the major fimbrial subunits were attributable to Lpf function, we complemented the Δ*lpfA1*, Δ*lpfA2*, and Δ*lpfA1*Δ*lpfA2* strains using plasmid-borne expression of the corresponding *lpf1* or *lpf2* operons. In epithelial cell adhesion assays, complementation restored adherence of the Δ*lpfA2* and double mutant strains to wild-type levels, while Δ*lpfA1* showed no strong baseline defect (Fig. S3). These data support the specificity of the phenotypes associated with disruption of the Lpf systems.

We next examined whether the contribution of Lpf to colonization was altered in a different intestinal environment using *sox10* mutant zebrafish, which have impaired intestinal motility (Kuhlman & Eisen, 2007). In wild-type fish, the Δ*lpfA1*Δ*lpfA2* strain showed a clear colonization defect relative to wild-type EHEC. In *sox10* animals, overall colonization was reduced and the difference between wild-type EHEC and the Δ*lpfA1*Δ*lpfA2* mutant was no longer apparent (Fig. 5D). Thus, the colonization advantage associated with Lpf depends on the host intestinal context, although the altered physiology of *sox10* animals precludes attributing this effect specifically to intestinal motility.

Together, these findings demonstrate an *in vivo* contribution of the Lpf fimbrial structure that does not require the distal tip adhesins under the conditions tested. This distinction parallels the requirement for the major fimbrial subunits, but not the tip adhesins, in surface-dependent regulatory feedback.

## DISCUSSION

Our findings identify a regulatory feedback circuit linking long polar fimbriae, surface sensing, and intestinal colonization. Lpf are established colonization factors whose expression responds to multiple host-associated conditions, but how these signals are integrated and whether Lpf themselves contribute to this regulation have remained unclear. Here, we show that *lpf1* and *lpf2* integrate metabolic and surface-associated signals through distinct promoter regions. Once expressed, Lpf promote surface-dependent activation of *ler* and *lpf*, establishing a positive-feedback circuit. Importantly, this response requires the major fimbrial subunits but not the distal tip adhesins, a distinction that is also observed during intestinal colonization.

The organization of the *lpf* promoters provides a mechanism for integrating different environmental inputs. Bile induced both long and short promoter constructs, indicating that bile responsiveness is retained within the proximal promoter regions. In contrast, removal of the distal regions markedly increased basal expression while largely eliminating induction following surface association. H-NS binding to the distal *lpf1* regulatory region has been demonstrated experimentally (Laaberki *et al*., 2006), and the corresponding *lpf2* region contains AT-rich sequences consistent with H-NS-dependent regulation. Thus, distal repression appears to establish the regulatory capacity for surface-dependent induction, whereas metabolic regulation can occur independently through the proximal promoter. The effects of bile and promoter truncation were qualitatively consistent across β-galactosidase, GFP, and qRT-PCR measurements, despite the greater dynamic range of the enzymatic β-galactosidase reporter. Together, these findings suggest that metabolic and surface-associated signals are integrated through separable features of the *lpf* promoters.

A central finding of this study is that Lpf themselves contribute to surface-dependent regulation. Loss of the major fimbrial subunits had little effect on basal *lpf* expression in planktonic cells but reduced induction following surface association. This phenotype was not attributable to impaired attachment, because cells were immobilized under conditions in which surface association does not depend on Lpf. Lpf therefore appear to enhance the regulatory response that follows surface contact rather than being required for initial surface association. This feedback intersects with the LEE regulatory network through Ler, a positive regulator of *lpf* expression (Torres *et al*., 2007a, Torres *et al*., 2008). Major fimbrial mutants similarly showed reduced surface-dependent *ler* activation without altered planktonic expression. While fimbrial deletion has been shown to cross-talk to regulation of other fimbriae in some cases (Gonyar *et al*., 2024), this does not appear to be the case here, since deletion mutants show unaltered expression in planktonic cultures (Fig. 3). Together, these findings support a reinforcing circuit in which Lpf structures are not essential for, but promote surface-dependent Ler activation, which can in turn enhance *lpf* expression. The pathway connecting Lpf to Ler remains to be determined.

An important feature of this response is its dependence on the major fimbrial subunits but not the distal tip adhesins. Chaperone–usher fimbriae are generally viewed as adhesive structures in which distal adhesins confer receptor specificity while the fimbrial shaft provides structural support. However, fimbrial shafts can also have important biomechanical properties, including the capacity to extend and dissipate force (Spaulding *et al*., 2018). Deletion of *lpfA1* and *lpfA2* impaired surface-dependent *lpf* and *ler* activation, whereas deletion of the distal tip adhesins LpfE1 or LpfD′2 did not. This distinction suggests that Lpf have a function during surface association that is separable from canonical tip-mediated adhesion.

The same distinction was observed *in vivo*. Loss of the major fimbrial subunits impaired intestinal colonization, whereas deletion of the distal tip adhesins did not significantly affect bacterial burden under the conditions tested. These findings do not imply that Lpf tip adhesins are generally dispensable for adhesion; Previous studies have demonstrated adhesive activities for Lpf at least in some contexts (Lloyd *et al*., 2012, Zhou *et al*., 2019, Torres *et al*., 2002, Jordan *et al*., 2004). Rather, our findings identify an additional function of Lpf during colonization that does not require the distal tip adhesins. Consistent with the importance of host context, the colonization difference between wild-type and Δ*lpfA1*Δ*lpfA2* bacteria was no longer apparent in *sox10* zebrafish. Because these animals have multiple intestinal abnormalities in addition to impaired motility, this result does not establish which host factor is responsible, but indicates that the contribution of Lpf is influenced by the intestinal environment.

Together, our findings support a model in which Lpf connect environmental sensing with transcriptional regulation during colonization. Metabolic and surface-associated signals are integrated through distinct features of the *lpf* promoters, while Lpf structures promote surface-dependent activation of Ler and their own expression. The requirement for the major fimbrial subunits, but not the distal tip adhesins, in both regulatory feedback and intestinal colonization reveals a role for Lpf beyond tip-mediated adhesion and expands the functions of fimbrial structures during host colonization.

## EXPERIMENTAL PROCEDURES

### Bacterial strains and growth conditions

Enterohemorrhagic *Escherichia coli* (EHEC) strain 86– 24 (O’Brien *et al*., 1983) was used as the wild-type background for all experiments, and all generated strains are listed in Table 1. In-frame deletion mutants (Δ*lpfA1*, Δ*lpfA2*, Δ*lpfA1*Δ*lpfA2*, Δ*lpfE1*, Δ*lpfD’2*) were generated in this background. Bacteria were routinely grown at 37°C in Luria–Bertani (LB) medium or Dulbecco’s Modified Eagle Medium (DMEM) without sodium carbonate, sodium pyruvate, L-glutamine, phenol red; supplemented with 0.5% sterile filtered glucose; adjusted to pH 6.5, under shaking or static conditions as indicated. Where specified, DMEM was supplemented with 0.15% sterile bile, as previously described (Arenas-Hernandez *et al*., 2014). Were indicated, antibiotics were added for plasmid maintenance and strain selection at the following concentrations: ampicillin (100 µg/ml), tetracycline (20 µg/ml), kanamycin (50 µg/ml), chloramphenicol (25 µg/ml).

**Table 1.**
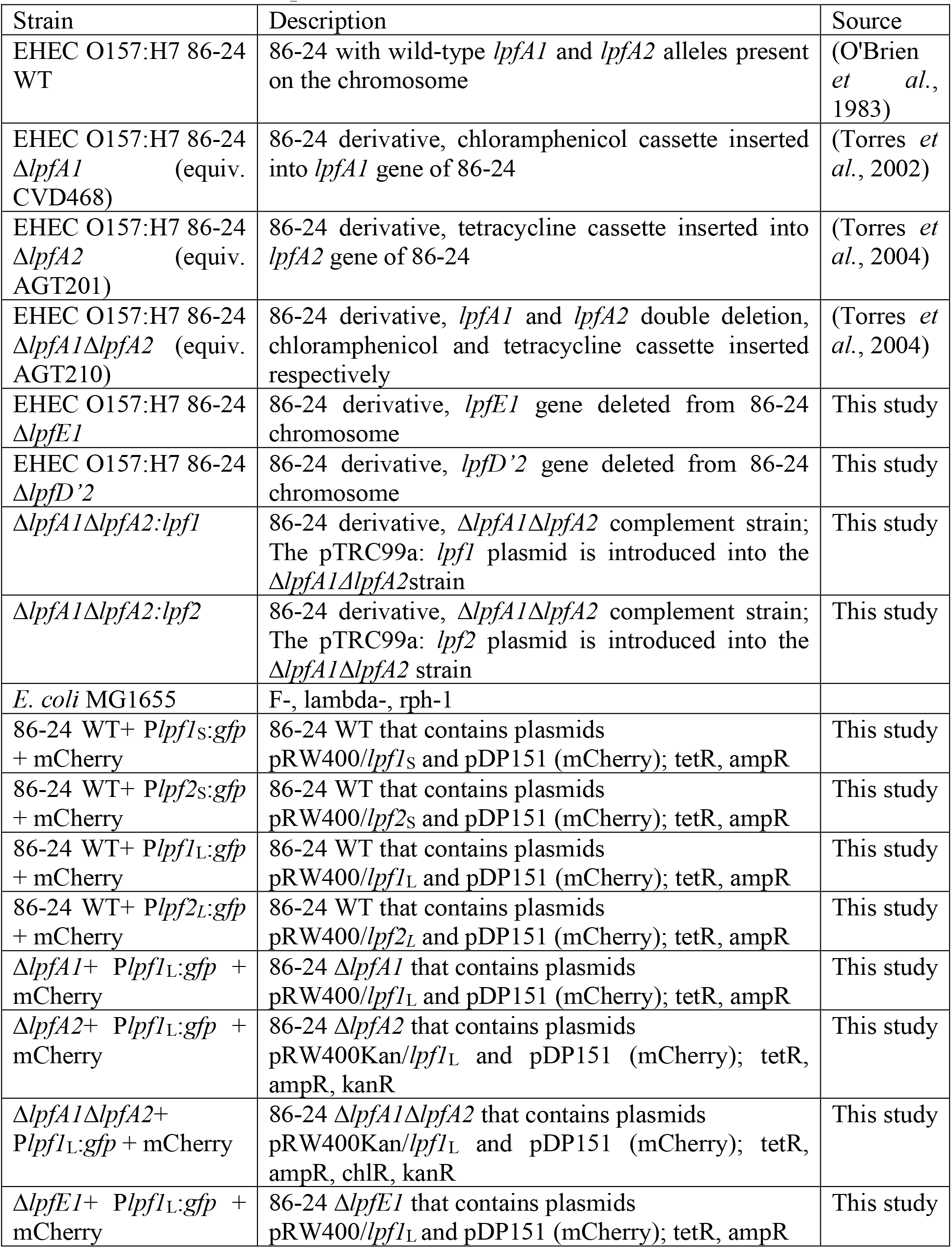
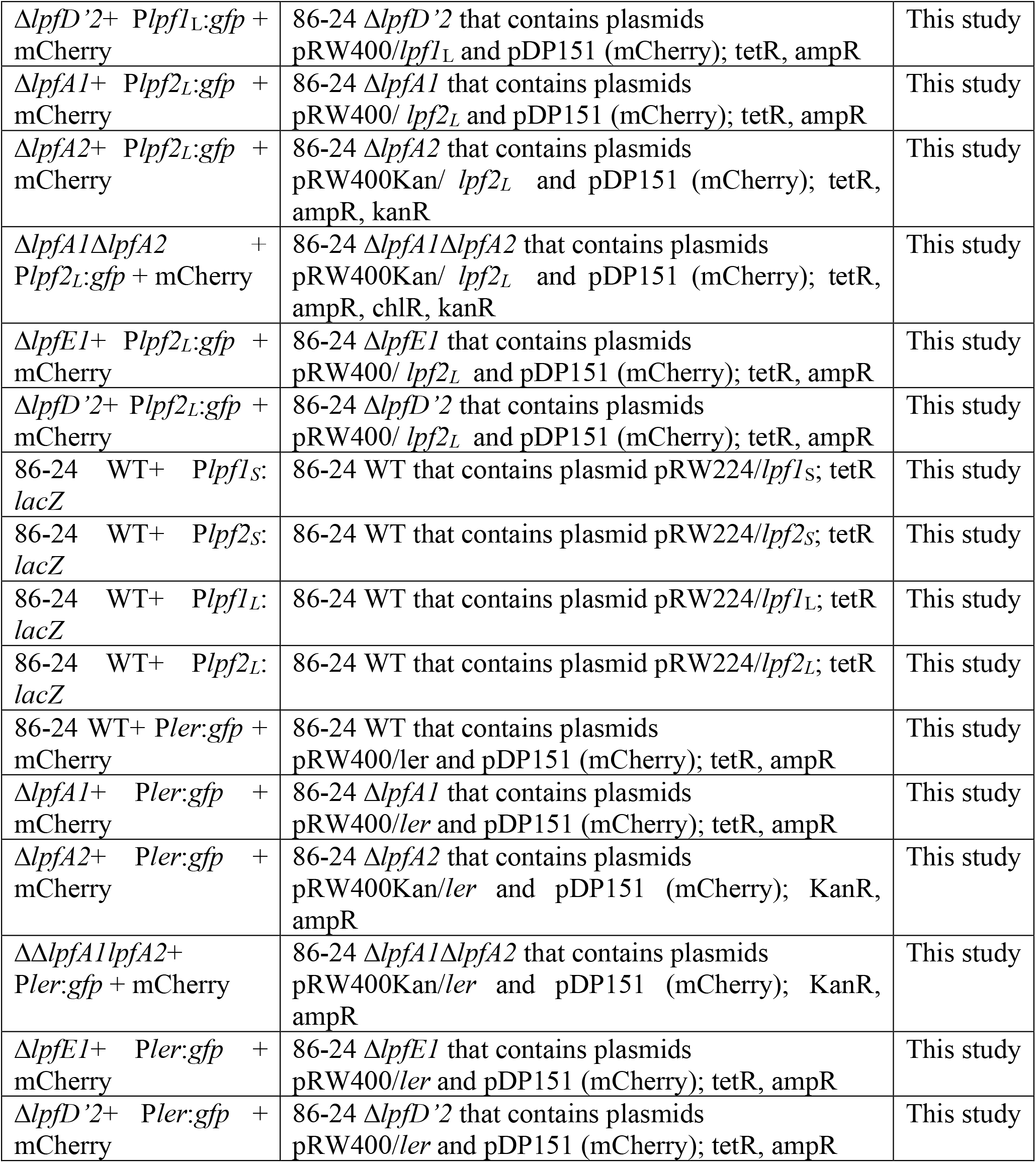
Strains used for experiments.

### Construction of promoter–reporter plasmids

Plasmids used in this study are listed in Table 2. Promoter regions upstream of *lpf1* and *lpf2* were amplified from EHEC strain 86-24 genomic DNA. Two promoter lengths were used: a long fragment (-350 bp for lpf1 and -310bp for lpf2 upstream of the translation start codon up to +10 bp) encompassing predicted regulatory regions including H-NS–associated binding sites, and a truncated fragment (-200 bp upstream for both lpf1 and lpf2) lacking distal upstream sequences. PCR products were generated and cloned upstream of either *lacZ* (into pRW224) or *gfp* (into pRW400) reporter genes using restriction enzyme cloning or Gibson assembly. Constructs were sequence-verified prior to use. Reporter plasmids were introduced into EHEC strains by electroporation.

**Table 2.**
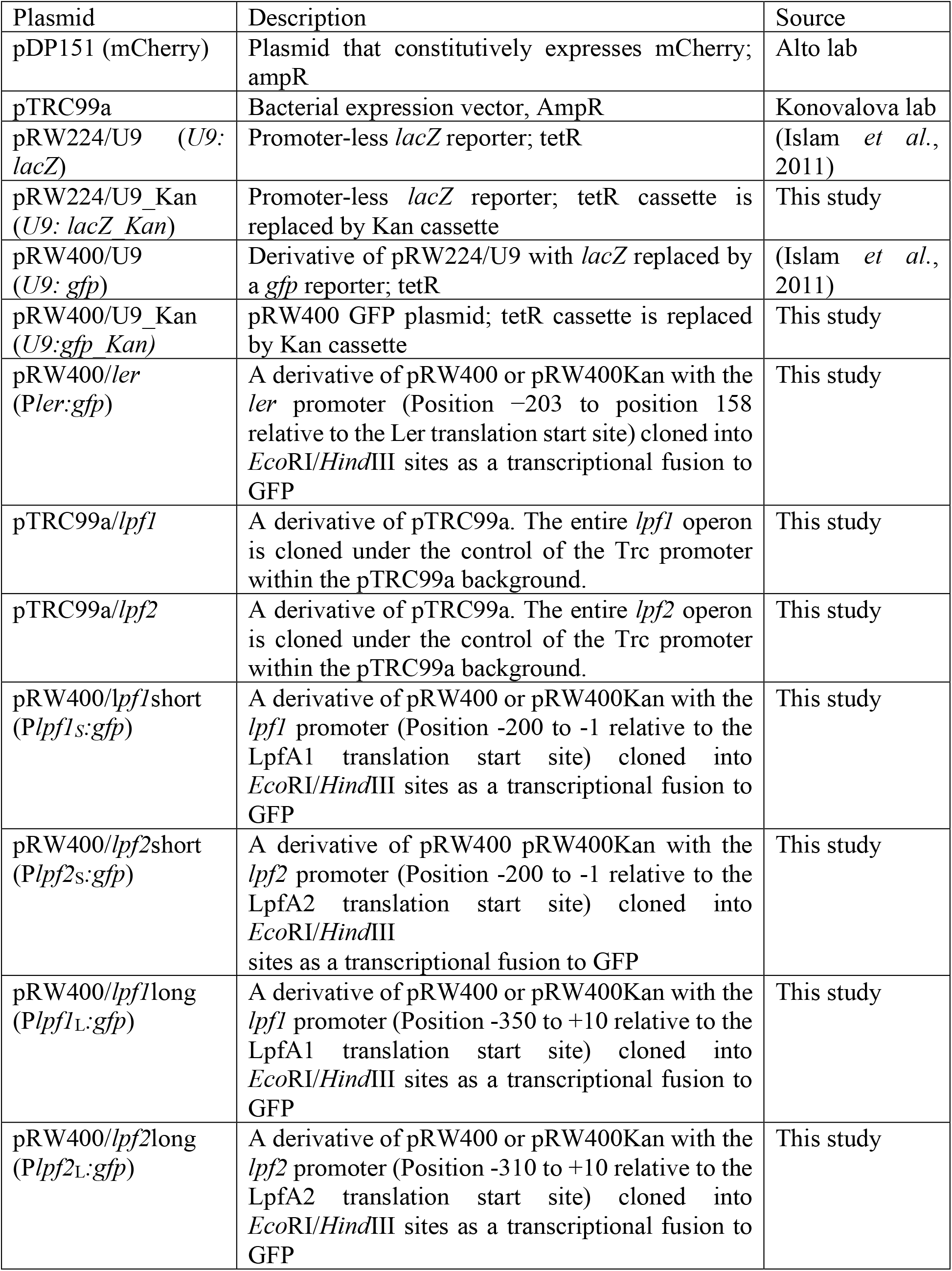
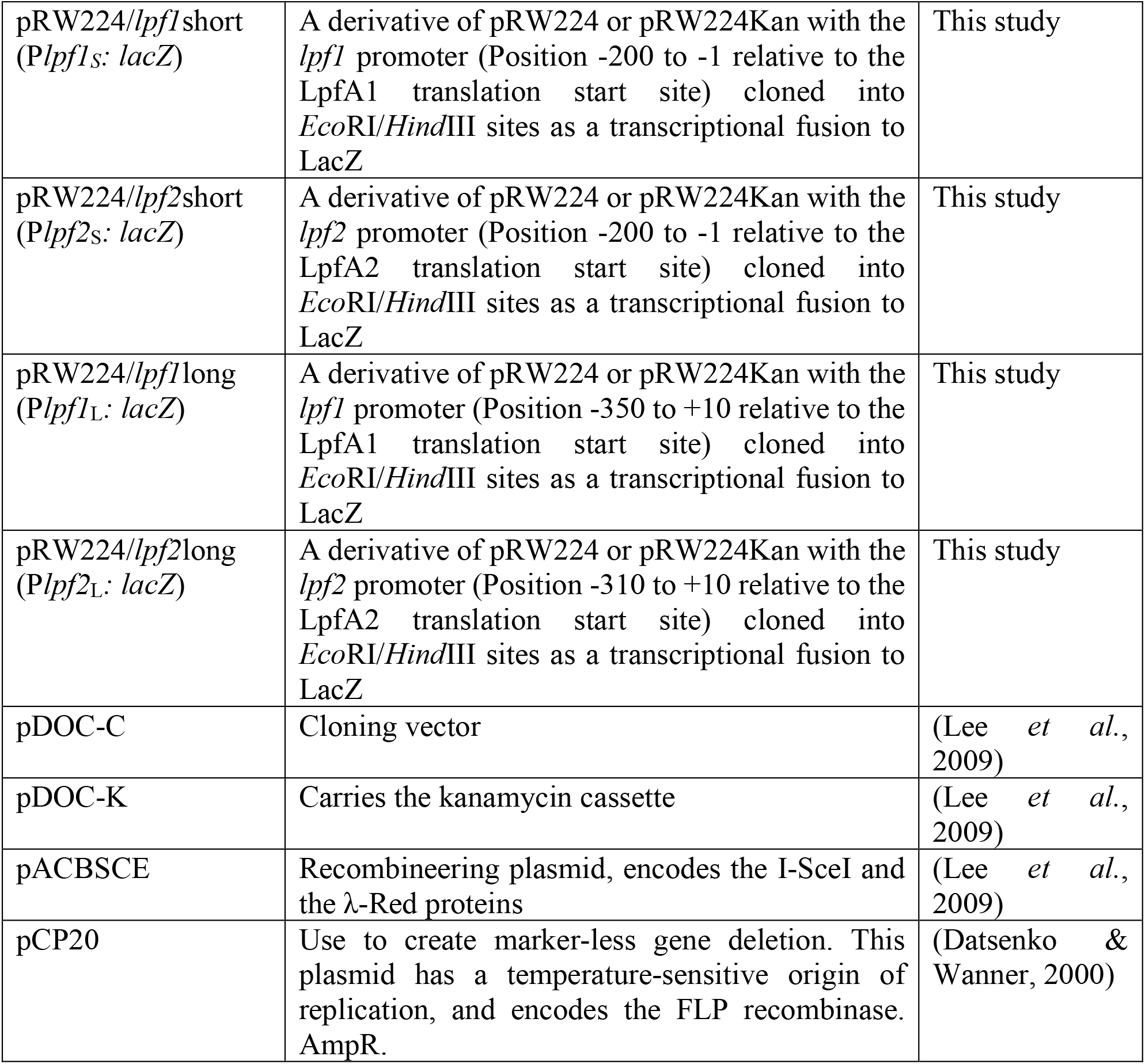
Plasmids used for experiments.

### Generation of deletion mutants

In-frame deletion mutants of *lpfA1, lpfA2*, and the double mutant were a gift from the Torres lab. Deletion mutants for the tip adhesin genes (*lpfE1, lpfD’2*) were constructed using gene doctoring as described previously (Lee *et al*., 2009). A Kan cassette was used for selection and removed where appropriate using FLP recombinase (pCP20) to generate markerless deletions. All mutants were confirmed by PCR and Sanger sequencing.

### Complementation plasmids and epithelial cell adhesion assays

For genetic complementation, the complete *lpf1* or *lpf2* operons were cloned into pTrc99A to generate pTrc-*lpf1* and pTrc-*lpf2* complementation plasmids. Constructs were sequence-verified and introduced into the Δ*lpfA1*Δ*A2* deletion strain by electroporation. As control, pTrc99A alone was introduced into wild type and Δ*lpfA1*Δ*A2* deletion strains. Functional complementation was assessed using HeLa cell adhesion assays. Monolayers (80% confluency) were infected with wild-type, mutant, or complemented strains in DMEM at an MOI of 30. After incubation for 5 hours at 37°C, non-adherent bacteria were removed by extensive washing with PBS. Cells were lysed with PBS containing 1% Triton X100, and adherent bacteria were quantified by serial dilution plating for CFU enumeration. Assays were performed in duplicate wells and at least three independent biological replicates.

### β-galactosidase assays

Promoter activity in planktonic cultures carrying lacZ reporters (Table 1, 2) was quantified using standard β-galactosidase (Miller) assays as described previously (Miller, 1972). Overnight cultures were diluted into fresh DMEM (without sodium carbonate, sodium pyruvate, L-glutamine, phenol red; supplemented with 0.5% sterile filtered glucose; adjusted to pH 6.5), and grown to the indicated growth phases. Where applicable, 0.15% bile was added to cultures at an OD_600_ of 0.2, and cultures grown for 3.5 hours to mid-log phase. Experiments were performed with at least three independent biological replicates.

### RT-qPCR

Overnight cultures were diluted into fresh DMEM. Where applicable, 0.15% bile was added to cultures at an OD_600_ of 0.2, and cultures grown for 3.5 hours to mid-log phase. Total RNA was purified from 10 ml of culture using PureLink RNA Mini Kit (Invitrogen). Total RNA was treated with RNAse-free DNAse Set from Qiagen before clean-up with RNA Clean & Concentrator kit (Zymo Research). Purified RNA was used for first-strand cDNA synthesis using the qScript Ultra SuperMix reverse transcriptase kit (Quantabio) on a CFX Real-Time PCR System (Bio-Rad). Fold enrichment was calculated using the 2^-ΔΔCT^ method, normalized to the constitutively expressed *gapA* control.

### Fluorescence microscopy and image analysis

For planktonic cells, fluorescence was quantified following fixation with 3.7% formaldehyde using fluorescence microscopy as described below. For validation experiments, fluorescence output was directly compared to β-galactosidase measurements across identical conditions. To quantify GFP in surface-associated cells, bacterial cultures were placed on sterile agarose pads and incubated at 37°C under static conditions for 1 hr. Imaging was performed using an Olympus FV3000 confocal microscope equipped with a 100x/1.40 NA oil-immersion objective and a GFP and mCherry filter set, at 2.5% laser power. The resulting images were processed in Olympus cellSens software and exported as TIFs. TIFs were analyzed using ImageJ/Fiji. Regions of interest corresponding to bacterial cells were defined using threshold-based segmentation of the red channel. Mean fluorescence intensity was measured for GFP and values shown are normalized to pixel area and background corrected. At least three independent replicates, and at least 150 cells/condition were analyzed.

### Zebrafish husbandry

All zebrafish experiments were conducted in accordance with the Guide for the Care and Use of Laboratory Animals and were approved by the Institutional Animal Welfare Committee of the University of Texas Health Science Center, Houston (protocol number AWC-25-0073). The zebrafish (*Danio rerio*) used in this study were wild-type AB and *sox10*^−/−^ larvae, which lack a functional enteric nervous system and exhibit impaired gut motility (Dutton *et al*., 2001). The *sox10* fish were maintained as a heterozygous breeding colony (*sox10*^*−/+*^), and following breeding, experimental larvae (*sox10*^*−/*−^) were identified by a lack of pigmentation and separated from the remainder of the clutch. Adult zebrafish were housed in a recirculating aquaculture system at the University of Texas Health Science Center at Houston Laboratory Animal Medicine and Care facility under standard conditions (28°C, pH 7.5, 14:10 h light:dark cycle). Fish were maintained at a density of up to five adult fish per liter of system water and fed GEMMA Micro (Skretting) twice daily, and embryos were obtained by natural spawning. Fertilized embryos were briefly surface-sterilized in 0.05% sodium hypochlorite for 30 s, rinsed three times in sterile embryo (E3) medium, and raised in petri dishes containing E3 (10 mM HEPES, 5 mM NaCl, 0.17 mM KCl, 0.4 mM CaCl_2_, 0.67 mM MgSO_4_, pH 7.4) at 28°C on a 14:10 h light:dark cycle. For experiments, zebrafish larvae were used at 6–7 days post-fertilization (dpf), corresponding to the developmental window when the intestine is fully formed and peristaltic motility is active (Ng *et al*., 2005).

### Zebrafish colonization assays

Larvae were infected at 6 days post fertilization using food-borne delivery of EHEC via the protozoan *Paramecium caudatum*, as described previously (Flores *et al*., 2019, Stones *et al*., 2017). Paramecia were maintained as described (Flores *et al*., 2019). Briefly, paramecia were co-cultured with bacteria for 2 h and thoroughly washed to remove extracellular EHEC. *E. coli*–loaded paramecia were counted using an automated cell counter (Life Technologies Countess II) and adjusted to a final concentration of 2 × 10^5^ paramecia/mL in E3 medium. Zebrafish larvae (6 dpf) were exposed to loaded paramecia for 2 h at 30°C in 6-well plates. Following the 2h exposure, larvae were washed in E3 and maintained in fresh E3 at 28°C. After 24 hours, individual larvae were euthanized in 1.6 mg/mL tricaine, washed, and homogenized for bacterial enumeration on CHROMagar O157 plates as described previously (Flores *et al*., 2023). Samples with no detectable bacterial burden were plotted as 1 for visualization and statistical analysis (limit of detection 24 CFU/fish).

### *F*luorescence imaging of colonized zebrafish and image-based burden quantification

Zebrafish larvae were colonized with EHEC mCherry strains as described above, and after 24 hours, larvae were washed in E3, euthanized with 1.6 mg/mL tricaine, washed again, and fixed in 3.7% formaldehyde. Larvae were then embedded in 1% low-melting-point agarose in a glass-bottom plate. Mounted larvae were imaged using an Olympus FV3000 confocal microscope with a 10x objective and 561nm excitation filter, with 1μm step-size image stacks across the entire width of the mid-intestine. Images were processed using Olympus cellSens software and total fluorescence intensity across the intestinal tract was calculated across all slices and background-corrected using an ROI outside the larvae. Images shown are Maximum intensity projections across the acquired z-stacks.

## Acknowledgements

We would like to thank A. Torres (UTMB) for sharing Δ*lpfA1*, Δ*lpfA2*, and Δ*lpfA1*Δ*lpfA2* strains. This work for funded by NIH grants R01AI132354 and R21AI199107 to AMK.

## Notes

**Conflict of interest disclosure:** The authors declare no conflicts of interest.

### Competing Interest Statement

The authors have declared no competing interest.

## REFERENCES

Alsharif, G., Ahmad, S., Islam, M.S., Shah, R., Busby, S.J., and Krachler, A.M. (2015) Host attachment and fluid shear are integrated into a mechanical signal regulating virulence in Escherichia coli O157:H7. Proceedings of the National Academy of Sciences of the United States of America 112: 5503–5508.

Arenas-Hernandez, M.M., Rojas-Lopez, M., Medrano-Lopez, A., Nunez-Reza, K.J., Puente, J.L., Martinez-Laguna, Y., and Torres, A.G. (2014) Environmental regulation of the long polar fimbriae 2 of enterohemorrhagic Escherichia coli O157:H7. FEMS microbiology letters 357: 105–114.

Barba, J., Bustamante, V.H., Flores-Valdez, M.A., Deng, W., Finlay, B.B., and Puente, J.L. (2005) A positive regulatory loop controls expression of the locus of enterocyte effacement-encoded regulators Ler and GrlA. Journal of bacteriology 187: 7918–7930.

Baumler, A.J., and Heffron, F. (1995) Identification and sequence analysis of lpfABCDE, a putative fimbrial operon of Salmonella typhimurium. Journal of bacteriology 177: 2087–2097.

Baumler, A.J., Tsolis, R.M., and Heffron, F. (1996) The lpf fimbrial operon mediates adhesion of Salmonella typhimurium to murine Peyer’s patches. Proceedings of the National Academy of Sciences of the United States of America 93: 279–283.

Chassaing, B., Etienne-Mesmin, L., Bonnet, R., and Darfeuille-Michaud, A. (2013) Bile salts induce long polar fimbriae expression favouring Crohn’s disease-associated adherent-invasive Escherichia coli interaction with Peyer’s patches. Environmental microbiology 15: 355–371.

Coppens, F., Iyyathurai, J., Ruer, S., Fioravanti, A., Taganna, J., Vereecke, L., De Greve, H., and Remaut, H. (2015) Structural and adhesive properties of the long polar fimbriae protein LpfD from adherent-invasive Escherichia coli. Acta Crystallogr D Biol Crystallogr 71: 1615–1626.

Cordonnier, C., Etienne-Mesmin, L., Thevenot, J., Rougeron, A., Renier, S., Chassaing, B., Darfeuille-Michaud, A., Barnich, N., Blanquet-Diot, S., and Livrelli, V. (2017) Enterohemorrhagic Escherichia coli pathogenesis: role of Long polar fimbriae in Peyer’s patches interactions. Scientific reports 7: 44655.

Datsenko, K.A., and Wanner, B.L. (2000) One-step inactivation of chromosomal genes in Escherichia coli K-12 using PCR products. Proceedings of the National Academy of Sciences of the United States of America 97: 6640–6645.

Dutta, S., Galdamez, W., Nguyen, A.T., Odem, M.A., Thompson, L., Bosserman, R.E., and Krachler, A.M. (2026) Intestinal motility shapes spatiotemporal patterns of enterohemorrhagic Escherichia coli colonization and virulence gene expression in the host gut. Infection and immunity 94: e0065725.

Dutton, K.A., Pauliny, A., Lopes, S.S., Elworthy, S., Carney, T.J., Rauch, J., Geisler, R., Haffter, P., and Kelsh, R.N. (2001) Zebrafish colourless encodes sox10 and specifies non-ectomesenchymal neural crest fates. Development 128: 4113–4125.

Farfan, M.J., Cantero, L., Vidal, R., Botkin, D.J., and Torres, A.G. (2011) Long polar fimbriae of enterohemorrhagic Escherichia coli O157:H7 bind to extracellular matrix proteins. Infection and immunity 79: 3744–3750.

Flores, E., Dutta, S., Bosserman, R., van Hoof, A., and Krachler, A.M. (2023) Colonization of larval zebrafish (Danio rerio) with adherent-invasive Escherichia coli prevents recovery of the intestinal mucosa from drug-induced enterocolitis. mSphere 8: e0051223.

Flores, E., Thompson, L., Sirisaengtaksin, N., Nguyen, A.T., Ballard, A., and Krachler, A.M. (2019) Using the Protozoan Paramecium caudatum as a Vehicle for Food-borne Infections in Zebrafish Larvae. Journal of visualized experiments : JoVE.

Gonyar, L.A., Sauder, A.B., Mortensen, L., Willsey, G.G., and Kendall, M.M. (2024) The yad and yeh fimbrial loci influence gene expression and virulence in enterohemorrhagic Escherichia coli O157:H7. mSphere 9: e0012424.

Hu, J., Ross, B.N., Cieza, R.J., and Torres, A.G. (2015) Finding Regulators Associated with the Expression of the Long Polar Fimbriae in Enteropathogenic Escherichia coli. Journal of bacteriology 197: 3658–3665.

Islam, M.S., Bingle, L.E., Pallen, M.J., and Busby, S.J. (2011) Organization of the LEE1 operon regulatory region of enterohaemorrhagic Escherichia coli O157:H7 and activation by GrlA. Molecular microbiology 79: 468–483.

Jordan, D.M., Cornick, N., Torres, A.G., Dean-Nystrom, E.A., Kaper, J.B., and Moon, H.W. (2004) Long polar fimbriae contribute to colonization by Escherichia coli O157:H7 in vivo. Infection and immunity 72: 6168–6171.

Kuhlman, J., and Eisen, J.S. (2007) Genetic screen for mutations affecting development and function of the enteric nervous system. Dev Dyn 236: 118–127.

Laaberki, M.H., Janabi, N., Oswald, E., and Repoila, F. (2006) Concert of regulators to switch on LEE expression in enterohemorrhagic Escherichia coli O157:H7: interplay between Ler, GrlA, HNS and RpoS. International journal of medical microbiology : IJMM 296: 197–210.

Lee, D.J., Bingle, L.E., Heurlier, K., Pallen, M.J., Penn, C.W., Busby, S.J., and Hobman, J.L. (2009) Gene doctoring: a method for recombineering in laboratory and pathogenic Escherichia coli strains. BMC microbiology 9: 252.

Lloyd, S.J., Ritchie, J.M., Rojas-Lopez, M., Blumentritt, C.A., Popov, V.L., Greenwich, J.L., Waldor, M.K., and Torres, A.G. (2012) A double, long polar fimbria mutant of Escherichia coli O157:H7 expresses Curli and exhibits reduced in vivo colonization. Infection and immunity 80: 914–920.

Miller, J.H., (1972) Experiments in Molecular Genetics. Cold Spring Harbor Laboratory Press., Cold Spring Harbor, NY: .

Newton, H.J., Sloan, J., Bennett-Wood, V., Adams, L.M., Robins-Browne, R.M., and Hartland, E.L. (2004) Contribution of long polar fimbriae to the virulence of rabbit-specific enteropathogenic Escherichia coli. Infection and immunity 72: 1230–1239.

Ng, A.N., de Jong-Curtain, T.A., Mawdsley, D.J., White, S.J., Shin, J., Appel, B., Dong, P.D., Stainier, D.Y., and Heath, J.K. (2005) Formation of the digestive system in zebrafish: III. Intestinal epithelium morphogenesis. Developmental biology 286: 114–135.

O’Brien, A.O., Lively, T.A., Chen, M.E., Rothman, S.W., and Formal, S.B. (1983) Escherichia coli O157:H7 strains associated with haemorrhagic colitis in the United States produce a Shigella dysenteriae 1 (SHIGA) like cytotoxin. Lancet 1: 702.

Rojas-Lopez, M., Arenas-Hernandez, M.M., Medrano-Lopez, A., Martinez de la Pena, C.F., Puente, J.L., Martinez-Laguna, Y., and Torres, A.G. (2011) Regulatory control of the Escherichia coli O157:H7 lpf1 operon by H-NS and Ler. Journal of bacteriology 193: 1622–1632.

Sirisaengtaksin, N., Odem, M.A., Bosserman, R.E., Flores, E.M., and Krachler, A.M. (2020) The E. coli transcription factor GrlA is regulated by subcellular compartmentalization and activated in response to mechanical stimuli. Proceedings of the National Academy of Sciences of the United States of America 117: 9519–9528.

Spaulding, C.N., Schreiber, H.L.t., Zheng, W., Dodson, K.W., Hazen, J.E., Conover, M.S., Wang, F., Svenmarker, P., Luna-Rico, A., Francetic, O., Andersson, M., Hultgren, S., and Egelman, E.H. (2018) Functional role of the type 1 pilus rod structure in mediating host-pathogen interactions. Elife 7.

Stones, D.H., Fehr, A., Voelz, K., and Krachler, A.M. Zebrafish (Danio rerio) as a vertebrate model to study colonization, pathogenesis and transmission of food-borne Escherichia coli O157. Under review.

Stones, D.H., Fehr, A.G.J., Thompson, L., Rocha, J., Perez-Soto, N., Madhavan, V.T.P., Voelz, K., and Krachler, A.M. (2017) Zebrafish (Danio rerio) as a Vertebrate Model Host To Study Colonization, Pathogenesis, and Transmission of Foodborne Escherichia coli O157. mSphere 2.

Torres, A.G., Blanco, M., Valenzuela, P., Slater, T.M., Patel, S.D., Dahbi, G., Lopez, C., Barriga, X.F., Blanco, J.E., Gomes, T.A., Vidal, R., and Blanco, J. (2009) Genes related to long polar fimbriae of pathogenic Escherichia coli strains as reliable markers to identify virulent isolates. Journal of clinical microbiology 47: 2442–2451.

Torres, A.G., Giron, J.A., Perna, N.T., Burland, V., Blattner, F.R., Avelino-Flores, F., and Kaper, J.B. (2002) Identification and characterization of lpfABCC’DE, a fimbrial operon of enterohemorrhagic Escherichia coli O157:H7. Infection and immunity 70: 5416–5427.

Torres, A.G., Kanack, K.J., Tutt, C.B., Popov, V., and Kaper, J.B. (2004) Characterization of the second long polar (LP) fimbriae of Escherichia coli O157:H7 and distribution of LP fimbriae in other pathogenic E. coli strains. FEMS microbiology letters 238: 333–344.

Torres, A.G., Lopez-Sanchez, G.N., Milflores-Flores, L., Patel, S.D., Rojas-Lopez, M., Martinez de la Pena, C.F., Arenas-Hernandez, M.M., and Martinez-Laguna, Y. (2007a) Ler and H-NS, regulators controlling expression of the long polar fimbriae of Escherichia coli O157:H7. Journal of bacteriology 189: 5916–5928.

Torres, A.G., Milflores-Flores, L., Garcia-Gallegos, J.G., Patel, S.D., Best, A., La Ragione, R.M., Martinez-Laguna, Y., and Woodward, M.J. (2007b) Environmental regulation and colonization attributes of the long polar fimbriae (LPF) of Escherichia coli O157:H7. International journal of medical microbiology : IJMM 297: 177–185.

Torres, A.G., Slater, T.M., Patel, S.D., Popov, V.L., and Arenas-Hernandez, M.M. (2008) Contribution of the Ler- and H-NS-regulated long polar fimbriae of Escherichia coli O157:H7 during binding to tissue-cultured cells. Infection and immunity 76: 5062–5071.

Vergara, A.F., Vidal, R.M., Torres, A.G., and Farfan, M.J. (2014) Long polar fimbriae participates in the induction of neutrophils transepithelial migration across intestinal cells infected with enterohemorrhagic E. coli O157:H7. Frontiers in cellular and infection microbiology 4: 185.

Zhou, M., Ding, X., Ma, F., Xu, Y., Zhang, J., Zhu, G., and Lu, Y. (2019) Long polar fimbriae contribute to pathogenic Escherichia coli infection to host cells. Applied microbiology and biotechnology 103: 7317–7324.

