## Supplemental file for "Surface sensing and metabolic signals drive feedback regulation of long polar fimbriae in enterohemorrhagic *Escherichia coli* O157"

**This file includes:**

Figures S1-S4

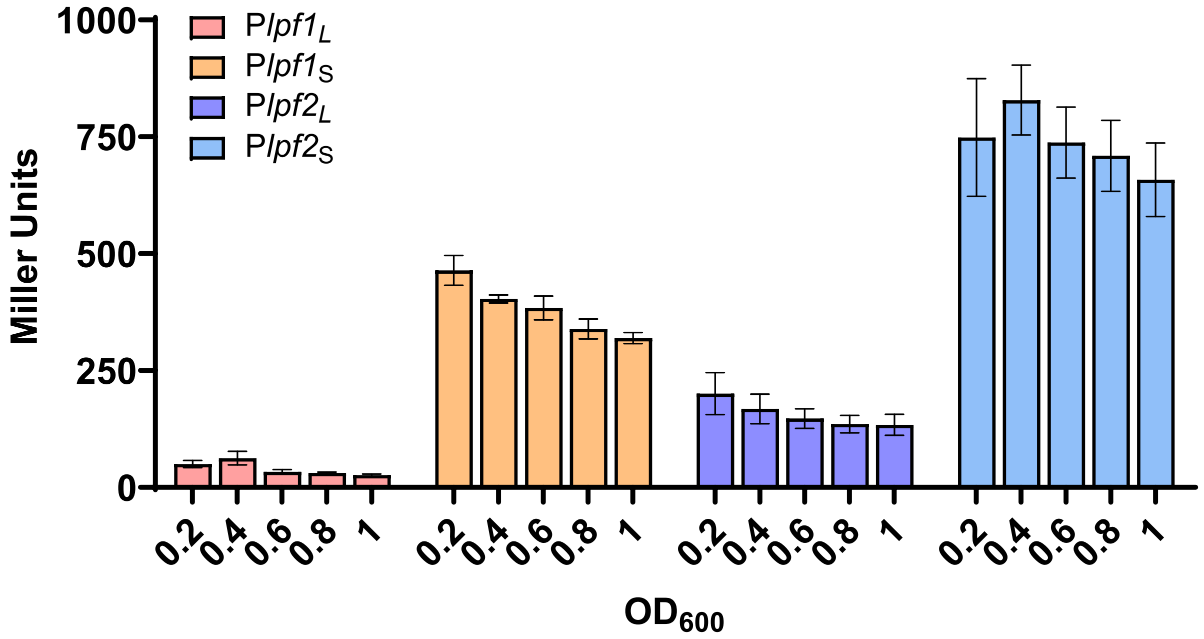


**Figure S1. Transcriptional activity of *lpf1* and *lpf2* promoters across growth phases.** β-galactosidase activity of *lpf1* and *lpf2* long and short promoter fusions to *lacZ* during growth in DMEM across growth phases ($OD_{600}$ of 0.2-1.0). Data represent mean ± SEM from three independent biological replicates;

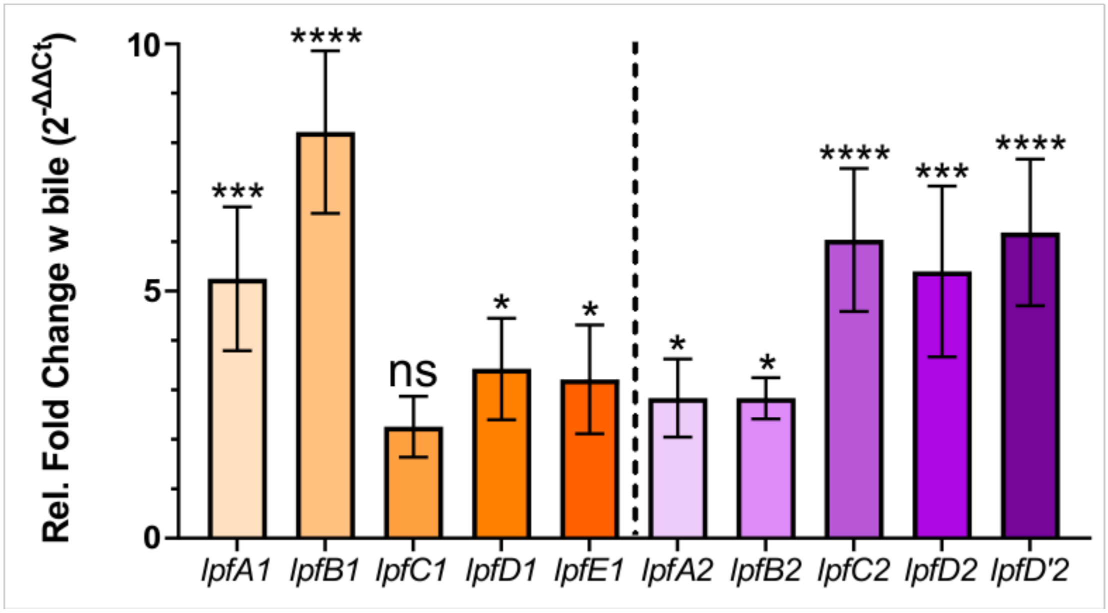


**Figure S2. Validation of bile-induced *lpf* expression by qRT-PCR.** Quantitative qRT–PCR analysis of *lpf1* and *lpf2* transcript levels following 0.15% bile exposure for 3.5 hours. Transcript levels were normalized using *gapA* as a housekeeping gene, and compared to those of cultures grown in DMEM without bile (1-fold). Data represent mean ± SEM from 4 independent biological replicates. Statistical significance was determined by lognormal one-way ANOVA with Dunnett's multiple-comparisons test (ns, $p \geq 0.05$; *$p < 0.05$; ***$p < 0.001$; ****$p < 0.0001$).

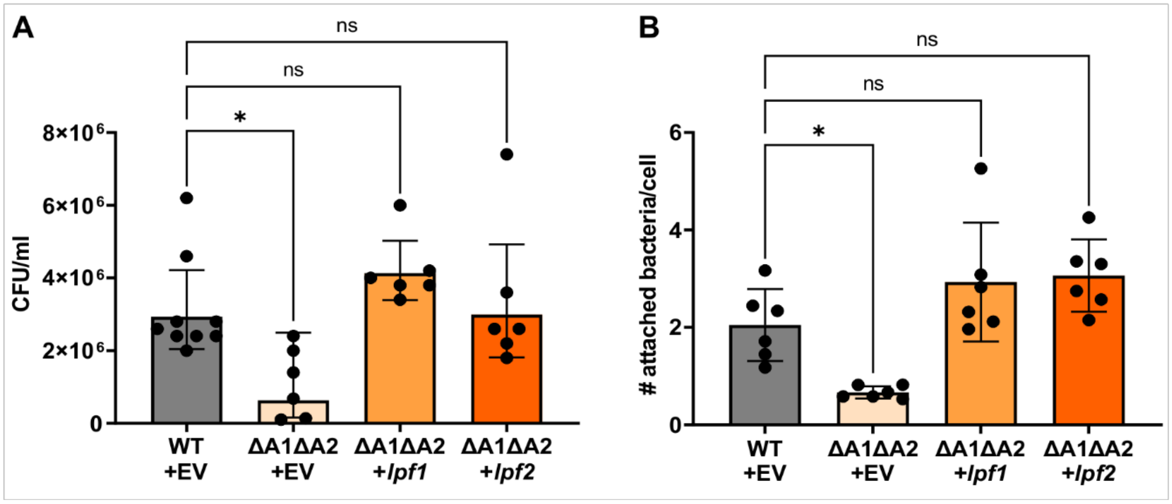


**Figure S3. Complementation of *lpf* deletion restores epithelial cell adhesion.** HeLa cell adhesion assays were performed to assess whether the Δ*lpfA1*Δ*lpfA2* mutant could be complemented in trans. Adherence of Δ*lpfA1*Δ*lpfA2* strain containing either pTrc empty vector (EV) or pTrc and the entire *lpf1* or *lpf2* operon, respectively, was compared to wild-type EHEC+EV as a positive control. Infected cell layers were washed and either **(A)** detergent lysed, and adherent EHEC quantified by dilution plating, or **(B)** fixed, stained, and adherent EHEC/Hela cell counted. Data represent mean ± SEM and individual data (n=6 for Δ*lpfA1*Δ*lpfA2* and 9 for WT in A; n=6 each in B). Statistical significance was determined by Borwn-Forsythe and Welch ANOVA tests, with Dunnett's T3 multiple comparisons test (ns, $p \geq 0.05$; $*p < 0.05$).

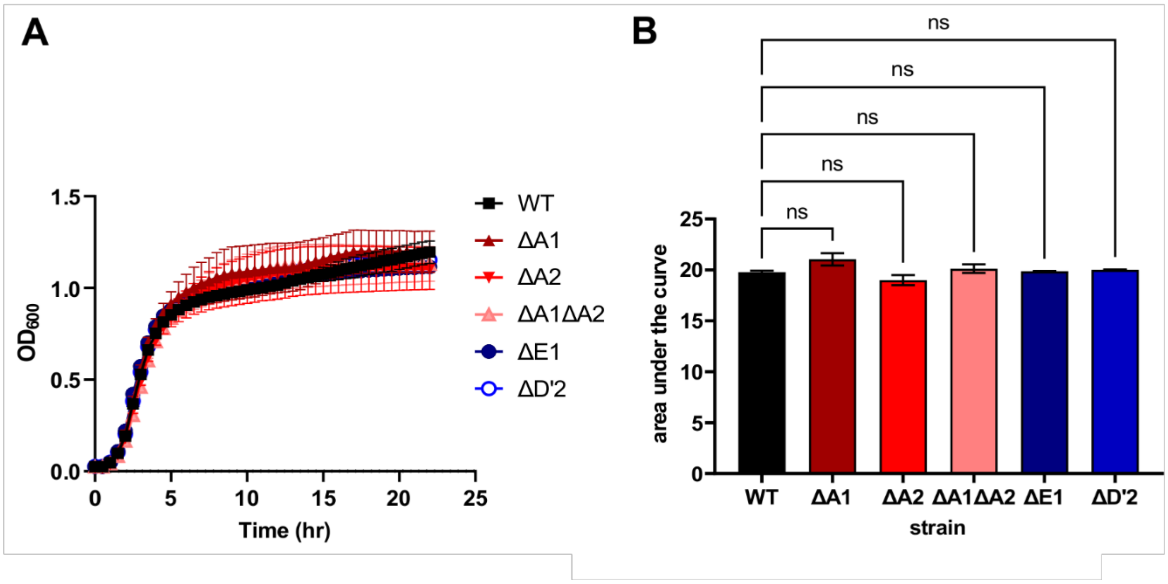


**Figure S4. Growth of wild-type EHEC and *lpf* deletion strains.** Strains were grown overnight and used to inoculate fresh cultures ($OD_{600}$ 0.025) in sterile 24well plates, covered with Breathe-easy gas permeable membranes, and grown at 37°C shaking. $OD_{600}$ was measured every 30 minutes for 22 hours **(A)**. **(B)** Data represent area under the curve for growth curves (mean ± SEM from 4 independent biological replicates). Statistical significance was determined by one-way ANOVA with Dunnett's multiple-comparisons test (ns, $p \geq 0.05$).
